# Optogenetic activation of afferent pathways in brain slices and modulation of responses by volatile anesthetics

**DOI:** 10.1101/2020.03.02.973115

**Authors:** Caitlin A. Murphy, Aeyal Raz, Matthew I. Banks

## Abstract

Anesthetics influence consciousness in part via their actions on thalamocortical circuits. However, the extent to which volatile anesthetics affect distinct cellular and network components of these circuits remains unclear. *Ex vivo* brain slices provide a means by which investigators may probe discrete components of complex networks and disentangle potential mechanisms underlying the effects of volatile anesthetics on evoked responses. To isolate potential cell type- and pathway-specific drug effects in brain slices, investigators must be able to independently activate afferent fiber pathways, identify non-overlapping populations of cells, and apply volatile anesthetics to tissue in aqueous solution. In this protocol, we describe methods to measure optogenetically-evoked responses to two independent afferent pathways to neocortex in *ex vivo* brain slices. We record extracellular responses to assay network activity and conduct targeted whole-cell patch clamp recordings in somatostatin- and parvalbumin-positive interneurons. We also describe a means by which to deliver physiologically relevant concentrations of isoflurane via artificial cerebral spinal fluid to modulate cellular and network responses.

**SUMMARY:** *Ex vivo* brain slices can be used to study the effects of volatile anesthetics on evoked responses to afferent inputs. We employ optogenetics to independently activate thalamocortical and corticocortical afferents to non-primary neocortex, and we modulate synaptic and network responses with isoflurane.

## INTRODUCTION

Volatile anesthetics have been used ubiquitously in a variety of clinical and academic settings for more than a century. Distinct classes of anesthetics have unique, often non-overlapping molecular targets^1-3^, yet nearly all of them produce unconsciousness. While their behavioral effects are quite predictable, the mechanisms by which anesthetics induce loss of consciousness are largely unknown. Anesthetics may ultimately influence both the level and contents of consciousness via actions on corticothalamic circuits, disrupting integration of information throughout the cortical hierarchy^4-9^. More broadly, modulation of corticothalamic circuits may play a role in experimentally^10^ or pharmacologically^11^ altered states of consciousness, and may also be implicated in sleep^12^ and in pathophysiological disorders of consciousness^13,14^.

The elusiveness of the mechanisms underlying loss and return of consciousness during anesthesia may be attributed partially to non-linear, synergistic actions of anesthetics at the cellular, network, and systems levels^15^. Isoflurane, for example, suppresses activity within select brain regions^16-18^, impairs connectivity between distant brain regions^19-23^, and diminishes synaptic responses in a pathway-specific manner ^24,25^. Which effects of anesthetics, from the molecular to the systems level, are necessary or sufficient to effect loss of consciousness remains unclear. In addition to substantive clinical investigations of consciousness using non-invasive techniques^19,20,26^, it is important that experimentalists seek to disentangle the distinct cellular and network interactions that subserve the conscious experience.

By simplifying the complex interactions found in the intact brain, *ex vivo* brain slices allow study of isolated components of the brain’s dynamic systems^9^. A reduced slice preparation combines the benefits of relatively intact anatomical structures of local neural circuits with the versatility of *in vitro* manipulations. However, until recently, methodological constraints have precluded study of synaptic and circuit properties of long-range inputs in brain slices^27,28^; the tortuous path of corticothalamic fiber tracts made activation of independent afferent pathways all but impossible by electrical stimulation, even for a careful geometer.

Investigating the effects of anesthetic agents on brain slice preparations presents additional challenges. Absent an intact respiratory and circulatory system, anesthetic agents must be bath-applied, and concentrations carefully matched to estimated effect site concentrations. For many intravenous anesthetic agents, the slow rate of equilibration in the tissue renders traditional pharmacological investigations laborious ^29,30^. Investigating the effects volatile gas anesthetics in *ex vivo* preparations is more tractable, but also presents challenges. These include converting inhaled partial pressure doses to aqueous concentrations, and the need for a modified delivery system of the drug to the tissue via artificial cerebral spinal fluid^31^.

Here, we describe the methods by which investigators may i) capitalize on the well-documented physicochemical properties of the volatile anesthetic isoflurane for drug delivery to *ex vivo* brain slices, ii) activate pathway- and layer-specific inputs to a cortical area of interest with high spatiotemporal resolution, and iii) conduct simultaneous laminar recordings and targeted patch clamp recordings from select populations of neurons. Combined, these procedures allow investigators to measure volatile anesthetic-induced changes in a number of observable electrophysiological response properties, from the synaptic to local network level.

## PROTOCOL

### 1. Breed mice to express fluorescent reporter protein in interneuron subpopulations

1.1. Pair homozyogous, Cre-dependent tdTomato male mouse with either SOM-Cre female or PV-Cre female mouse. Other specific neuronal populations may be targeted by using the appropriate Cre lines, but here we will concentrate on the lines we chose to study.

1.2. Allow heterozygous offspring to mature to at least 3 weeks of age before proceeding.

### 2. Perform stereotaxic injection of viral construct

2.1. Adjust settings of micropipette puller for injection pipettes as indicated in instrument user manual (see Table 2 for recommended settings). Polish the ends of glass tubing and pull glass micropipette.

**Table 1.** Composition of artificial cerebral spinal fluid and intracellular solution. Reagents and concentrations for slicing ACSF (sACSF), experimental ACSF (eACSF), and intracellular pipette solution for patch clamp recordings are listed.

|  | <u>Slicing ACSF, sACSF (in mM)</u> | <u>Experiment ACSF, eACSF (in mM)</u> |
| --- | --- | --- |
| NaCl | 111 | 111 |
| NaHCO<sub>3</sub> | 35 | 35 |
| HEPES | 20 | 20 |
| KCl | 1.8 | 1.8 |
| CaCl<sub>2</sub> | 1.05 | 2.1 |
| MgSO<sub>4</sub> | 2.8 | 1.4 |
| KH<sub>2</sub>PO<sub>4</sub> | 1.2 | 1.2 |
| glucose | 10 | 10 |

|  | <u>Internal Solution</u> |
| --- | --- |
| K-gluconate | 140 |
| NaCl | 10 |
| HEPES | 10 |
| EGTA | 0.1 |
| MgATP | 4 |
| NaGTP | 0.3 |
|  | pH = 7.2 |

**Table 2.**
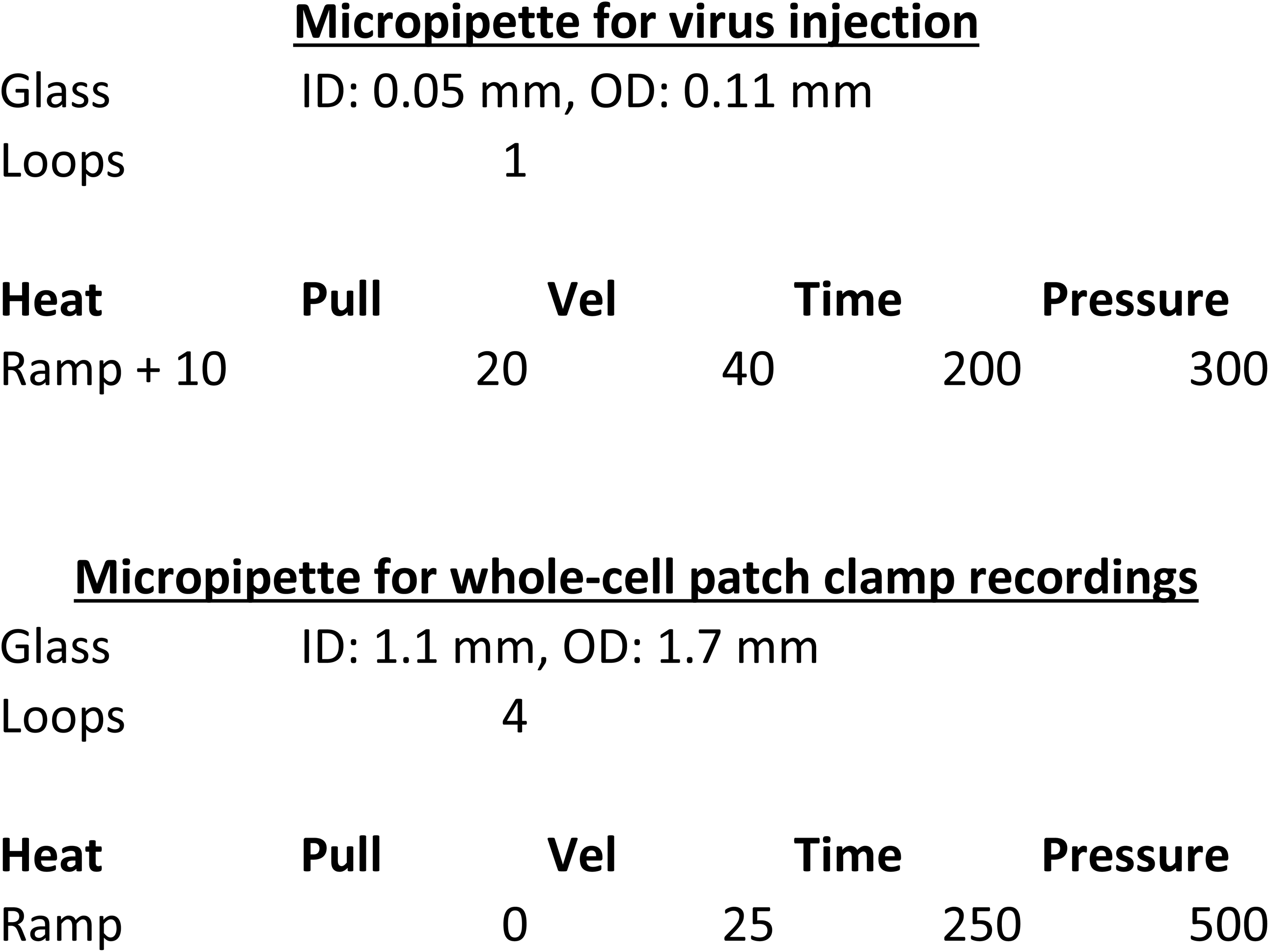
Recommended glass and parameters for pulling micropipettes for viral injections and whole-cell patch clamp recordings. Glass used for viral injection and whole-cell patch clamp recordings is described, as well as the parameters for pulling micropipettes using the Flaming/Brown micropipette puller by Sutter. Investigators should consult instruction manuals for micropipette puller for further recommendations or fine-tuning of settings.

2.2. Break the tip of the sharp end of the pipette. Pull 2-3 µL of non-compressable mineral oil up into pipette, followed by at least 1.0 µL of viral construct. The recombinant adeno-associated viral vector used in these experiments was AAV2-hSyn-hChR2(H134R)-EYFP.

2.3 Arrange sterile drape in surgical area. Sterilize tools for stereotaxic procedure and place on drape.

2.4 Anesthetize SOM-tdTomato or PV-tdTomato heterozygous animal using isoflurane (3% for induction, 1.5-2% for maintenance) and oxygen mixture. Periodically confirm surgical level of anesthesia with toe pinch throughout surgery.

2.5. Shave the top of the animal’s head. Apply 70% isopropyl alcohol and Betadine solution liberally to surgical area and ophthalmic ointment to eye sockets to prevent drying of membrane. Administer bupivacaine/lidocaine (1:1 ratio, 1.0 mg/kg) subcutaneously to surgical site for local anesthetic.

2.6. Fit animal into stereotaxic frame.

2.7. Use scalpel to make incision along sagittal axis of skin overlying the dorsal surface of the skull. Retract skin using forceps. Apply hydrogen peroxide and saline to skull.

2.8. On the surface of the skull, lightly mark the intersection of anterior and lateral coordinates with a cross in pencil. Drill a hole at the appropriate coordinates in the transverse plane (for cingulate cortex (Cg) injection: anterior 0.2 mm, lateral 0.3 mm; for posterior thalamus (Po) injection: posterior 2.25 mm, lateral 3.4 mm). (NOTE: Markings should extend beyond the boundaries of the burr hole to provide guidance for accurate placement of pipette.)

2.9. Re-position dorso-ventral arm of stereotaxic frame at 0° for injections into Cg, or 45° in the coronal plane for injections into Po.

2.10. Mount loaded injection pipette into microsyringe pump and attach syringe pump to dorso-ventral arm of the stereotaxic frame. Attach the microsyringe pump controller to the syringe pump.

2.11. Navigate pipette tip near (but not touching) the surface of the brain, at the intersection of markings created in Step 2.8. Carefully advance the pipette along its longitudinal axis into the brain either 0.9 mm (injection in Cg) or 3.1 mm (injection in Po). Wait 10 minutes before proceeding.

2.12. Inject 100 nL/min over 10 minutes. If welling of virus from pipette insertion site is observed, slow injection rate to 50 nL/min. Inject 1.0 µL of viral construct total.

2.13. After injection, wait 10 minutes before slowly retracting injection pipette.

2.14. Suture scalp incision closed and administer 2-5 mg/kg meloxicam subcutaneously.

2.15. Discontinue isoflurane and monitor animal during emergence from anesthesia. Allow to recover according to procedures described by institution’s Animal Care and Use Committee, including further administration of analgesics.

### 3. Prepare brain acute brain slices

3.1. Allow at least 3 weeks for expression of viral construct before harvesting tissue.

3.2. Prepare 1 L of artificial cerebral spinal fluid for slicing procedure (sACSF). See Table 1 for ingredients.

3.3. Throughout slicing procedure, supply sACSF with dissolved 95% O_2_/5% CO_2_ mixture, delivered via gas dispersion tube.

3.5. Prepare ice cold bath for vibrating blade microtome. Mount ice-cold specimen stage onto microtome and fix sapphire blade in place for tissue sectioning.

3.4. Anesthetize mouse with 3% isoflurane and oxygen until loss of righting reflex.

3.5. Decapitate mouse using guillotine and immediately submerge head in 4°C sACSF. To preserve the health of the tissue, the following steps should be completed as swiftly as possible.

3.6. Open skull cavity by making a small incision at the base of the skull and gently removing each skull plate. Gently remove underlying dura mater.

3.7. While the brain is still in the skull cavity, use the razor blade to remove the cerebellum. Make a second vertical cut along the sagittal plane in the left hemisphere, just lateral to the midline.

3.8. Prepare tissue block for sectioning.

3.8.1. Gently lift the brain from the skull cavity. Place the brain on the filter paper with the flat, sagittal plane down. Guide the filter paper over the blocking template and align the brain to the underlying template outline (Supplementary Figure 1).

**Figure 1.**
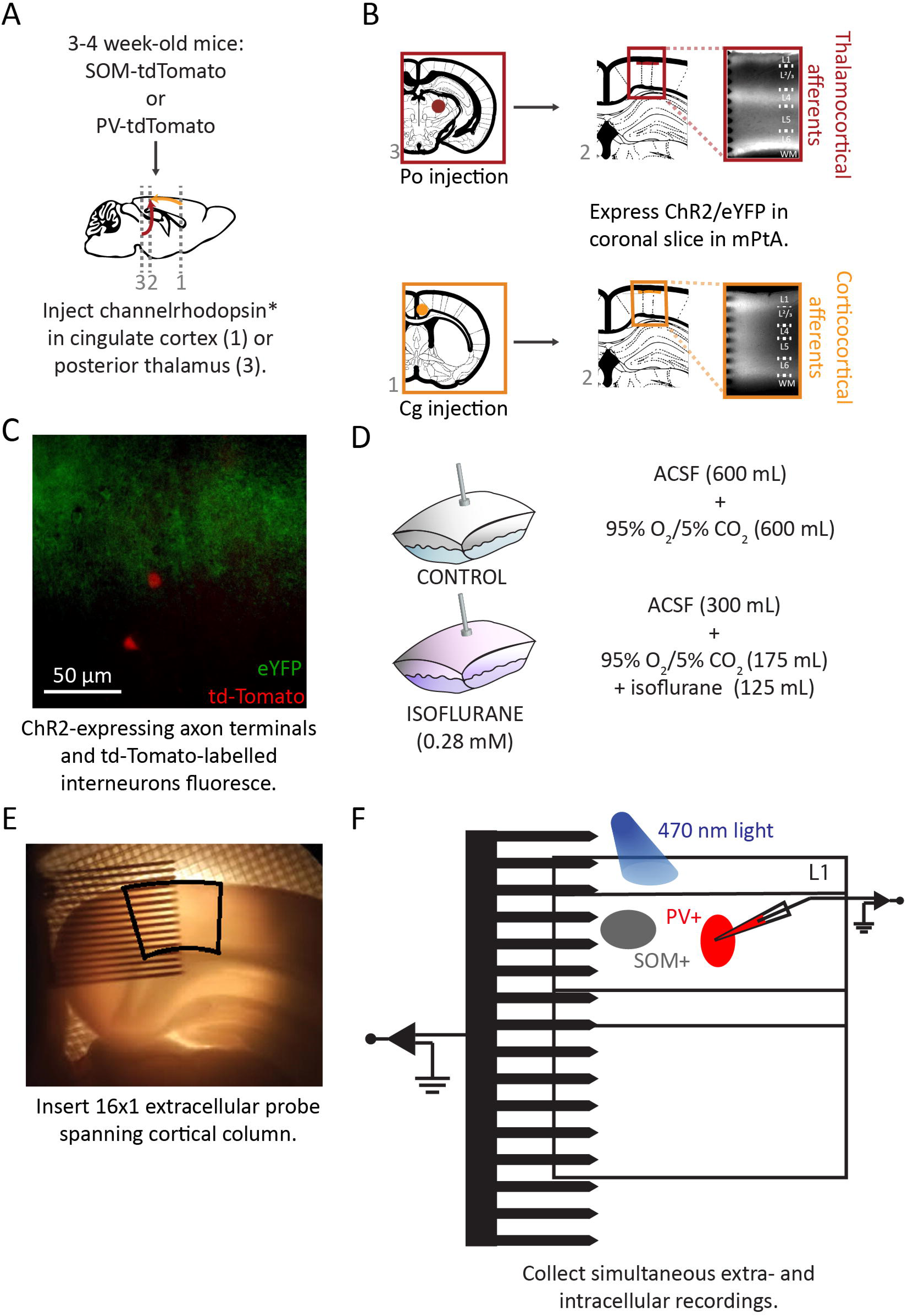
Injection of viral vector and preparation *ex vivo* coronal brain slices. (**A**) Schematic representation of injection of viral vector into SOM-tdTomato or PV-tdTomato hybrid mice. (**B**) Coronal slices of the medial parietal association area (mPtA) were harvested, and thalamocortical (top) or corticocortical (bottom) afferent fibers were identified by their eYFP reporter in layer 1. (**C**) Overlay of eYFP-labelled axon terminals in layer 1 (green) and tdTomato-labelled SOM+ interneurons (red) in superficial layer 2/3. (**D**) Sealed Teflon bags were prepared with a 50:50 solution-to-gas mixture. (**E**) Placement of a 16×1 probe into mPtA (black outline). (**F**) Schematic of the recording set-up in the cortical slice.

3.8.2. Make two parallel cuts in the coronal plane as indicated by the lines on the template. Add a small drop of sACSF to keep filter paper wet, if necessary.

3.8.3. Place tissue block in 4°C sACSF.

3.8.4. Apply a small amount of super glue to the ice-cold specimen stage.

3.8.5. Lift tissue block from cold sACSF, and use the corner of a Wypall towel to wick away excess sACSF. Glue the posterior coronal plane of the tissue block to the specimen stage, with the dorsal surface of the brain facing the sapphire blade.

3.9. Collect 500 µm-thick coronal brain slices. Place slices of interest on nylon mesh (Supplementary Figure 2) in 34°C sACSF and allow the container to reach room temperature. NOTE: For experiments described here, electrophysiological recordings were collected from a coronal section centered approximately 2.25 mm posterior to bregma to study a non-primary sensory area, medial secondary visual cortex (V2MM).

**Figure 2.**
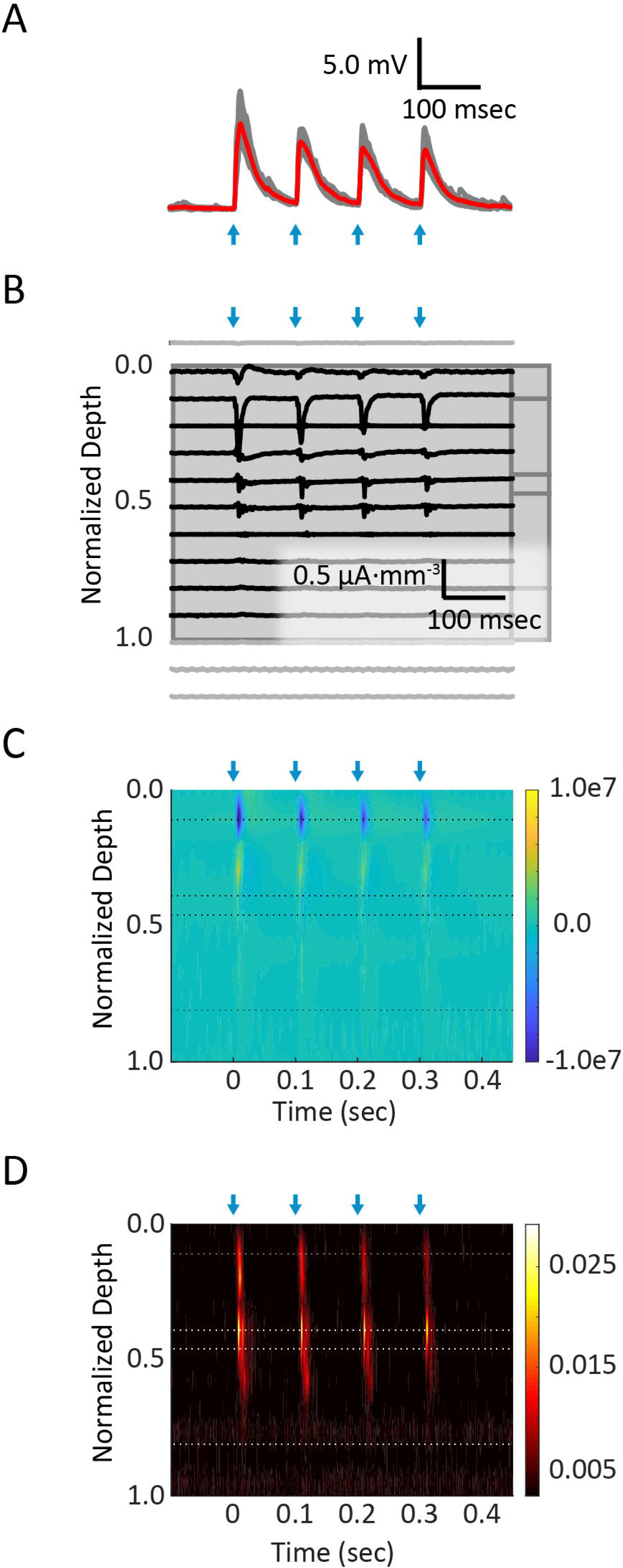
Simultaneous intracellular and multi-channel extracellular recordings in cortical slice. (**A**) Whole-cell current clamp patch recording from the soma of a layer 2/3 PV+ interneuron. Four pulses (2 msec each, blue arrows) of blue light (2.2 mW) at 10Hz were delivered to corticocortical axon terminals in L1. Average (red trace) of ten trials (grey traces) are shown. (**B**) Raw data from 16 channels of extracellular 16×1 probe. Channels placed in cortical tissue are shown in black, and those lying outside of cortex in grey. (C) A current source density diagram, extracted from the local field potential signal, shows synaptic current sinks (blue) in layer 1. (**D**) Multi-unit activity, generated by applying a high-pass filter to the local field potential signal, isolates spiking activity evoked in lower layers.

### 4. Prepare bags of eACSF containing dissolved volatile anesthetic isoflurane

4.1. Prepare 300 mL of a stock mixture of 3.0% isoflurane.

4.1.1. In a sealed Teflon gas bag, add ∼100 mL 95% O_2_/5% CO_2_ gas mixture to 20-30 mL of liquid isoflurane and a small amount of 0.9% saline. Wait at least 30 minutes to allow equilibration of isoflurane between liquid and gas phases.

4.1.2. In Determine the amount of saturated isoflurane gas, V_sat_, to add to the stock bag using the following equation:

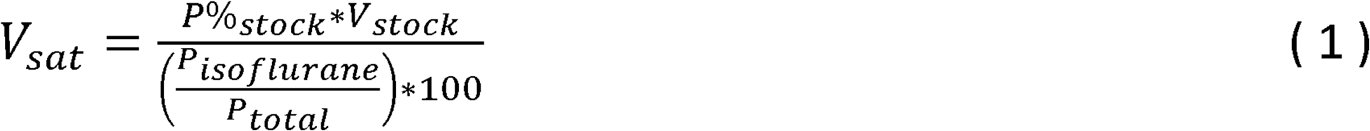

where P%_stock_ is the target composition of the stock gas (3% in this case), V_stock_ is the final volume of the stock gas bag, P_isoflurane_ is the partial pressure of isoflurane at room temperature (∼240 mmHg), and P_total_ is the atmospheric pressure (∼760 mmHg).

4.1.3. In Add the calculated amount of saturated gas to an empty gas bag, and fill the bag with a volume of 95% O_2_/5% CO_2_ gas mixture to bring total volume of stock bag to 300 mL.

4.2. In Prepare 2 L of artificial cerebral spinal fluid for perfusion of the slice during the experiment (eACSF). See Table 1 for ingredients. Dissolve 95% O_2_/5% CO_2_ gas mixture into solution.

4.3. Prepare two separate bags of CONTROL and ISOFLURANE solutions.

4.3.1. To an empty teflon gas bag, add 600 mL eACSF and 600 mL of 95% O_2_/5% CO_2_ gas mixture. Label this bag CONTROL.

4.3.2. To an empty teflon gas bag, add 300 mL eACSF. Label this bag ISOFLURANE.

4.3.3. Choose a physiologically relevant equilibrated gas phase concentration of isoflurane. Experiments were conducted using gas concentrations equivalent to 1.3% isoflurane. (Mice lose righting reflex, and presumably consciousness, at 0.9% inhaled isoflurane.)

4.3.4. Use the following equation to calculate the equivalent gas phase concentration at room temperature, P%(T _room_) ^31^:

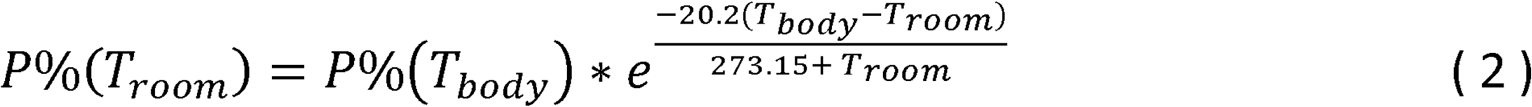

where P%(T_body_) is the physiologically relevant gas phase concentration chosen in Step 4.3.3, T_room_ is 25 °C, and T_body_ is 37 °C.

4.3.5. Use the following equation to determine volume of gas from stock gas bag, V_stock_, to add to the ISOFLURANE solution.

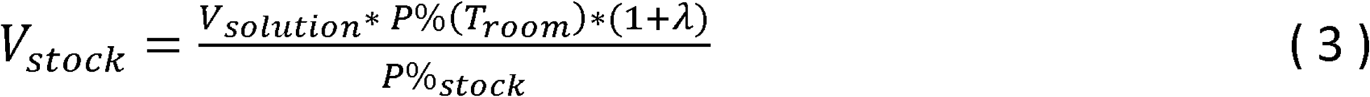

where V_solution_ is the volume of eACSF in ISOFLURANE bag (300 mL), P%(T_room_) is entered from equation (2), λ is the saline/gas Ostwald partition coefficient of isoflurane (λ = 1.2^32^), and P%_stock_ is the gas phase concentration of the stock gas bag (P%_stock_ = 3.0%).

4.3.6. To the ISOFLURANE solution bag, add the calculated volume of gas from the stock gas bag.

4.3.7. To the ISOFLURANE solution bag, add 95% O_2_/5% CO_2_ gas mixture until the total volume of gas is 300 mL.

4.4. Shake both CONTROL and ISOFLURANE bags on shaker for at least 1 hour to allow isoflurane phase equilibration.

4.5. After all data has been collected, you may verify the correct concentration by using an anesthetic gas monitor to measure equilibrated gas concentration of isoflurane above the remaining solution in bag.

4.6. Report experimental concentrations of volatile gases in aqueous units, as millimolar concentrations are more robust to changes in temperature. Use the following equation to convert room temperature gas phase concentration, P%(T_room_), to equivalent aqueous concentration (C aqueous, in mM) ^31^:

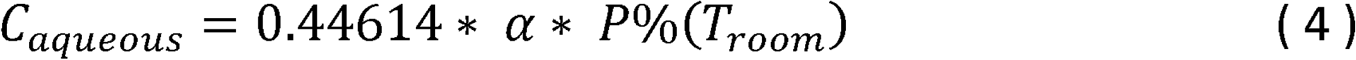

where α is the saline/gas Bunsen partition coefficient for isoflurane at 25°C^32^.

### 5. Prepare hardware and software for multi-channel recordings

5.1. Set up 16-channel data acquisition system according to manufacturer instructions.

NOTE: Several commercially available amplifiers and data acquisition systems can be used to collect multi-channel recordings. In the experiments described here, analog signals are delivered via an electrode reference panel to two Lynx-8 amplifiers, where they are amplified (2000x) and filtered (0.1-10kHz). Analog inputs to the data acquisition system (Digidata 1440A) are digitized at 40kHz.

5.2. Fasten the appropriate 16-channel headstage adaptor to a microscope micromanipulator. Orient the adaptor such that the female connector ports are facing downward.

5.3. Adjust the angle of operation of this micromanipulator such that it is oriented downward toward the recording chamber, at an angle approximately 70° relative to horizontal.

5.4. Connect the headstage input to a 16×1 NeuroNexus probe for *in vitro* electrophysiology via the headstage adaptor anchored to the micromanipulator.

5.5. Connect the headstage output connector to the data acquisition system.

5.6. Install appropriate software for data acquisition (e.g., Clampex 10). Configure 15 input channels to correspond to input signals from the first 15 multi-channel probe contacts. Configure the remaining channel to receive input from the intracellular electrode.

NOTE: Take care to consider electrode and adaptor maps when collecting and analyzing data, to ensure the appropriate signal corresponds to the electrode contact from which it was collected.

### 6. Configure light stimulation protocols

6.1. Set up Polygon400 or comparable light delivery device. Install PolyScan2 software.

6.2. Open PolyScan2 software. Choose hardware wiring configuration in which a Trigger Source (Digital/TTL Out) provides TRIGGER IN signal to the Polygon400, and the Polygon400 provides TRIGGER OUT signal to 470 nm LED.

6.3. Mount high-power objective lens. Using digital camera, calibrate high-power objective following on-screen instructions.

6.4. Create new profile sequence of light stimulation profiles.

6.4.1. Open the Profile Sequence Editor. Open a New Profile.

6.4.2. In the Pattern Editor, create a pattern of choice. In the experiments described here, a circle of diameter 150 µm is used to allow layer-specific activation of axon terminals.

6.4.3. To construct a profile sequence, copy and paste this profile for any number of trials.

6.4.4. Create a Waveform List that contains waveforms of any light intensity, pulse duration, or pulse number.

6.4.5. Randomly assign waveforms to each profile. Each profile with its assigned waveform corresponds to one trigger pulse from a Digital TTL input, or one trial.

6.4.6. Save the profile sequence.

6.5. In the data acquisition software, create a new protocol.

6.5.1. Set the number of trials to equal the number of profiles in the profile sequence just created in PolyScan2.

6.5.2. Choose signal inputs to match those configured in Step 5.6. Configure a protocol that provides a single digital TTL output, recording from these 16 input channels for an appropriate amount of time before and after the digital trigger.

### 7. Place multi-channel probe in ex vivo brain tissue slice

7.1. Perfuse bubbled eACSF (not in sealed bags) at 3-6 mL/min.

7.2. Transfer brain slice containing area of interest onto mesh grid in microscope perfusion chamber. Anchor with platinum harp (see Supplementary Figure 3).

**Figure 3.**
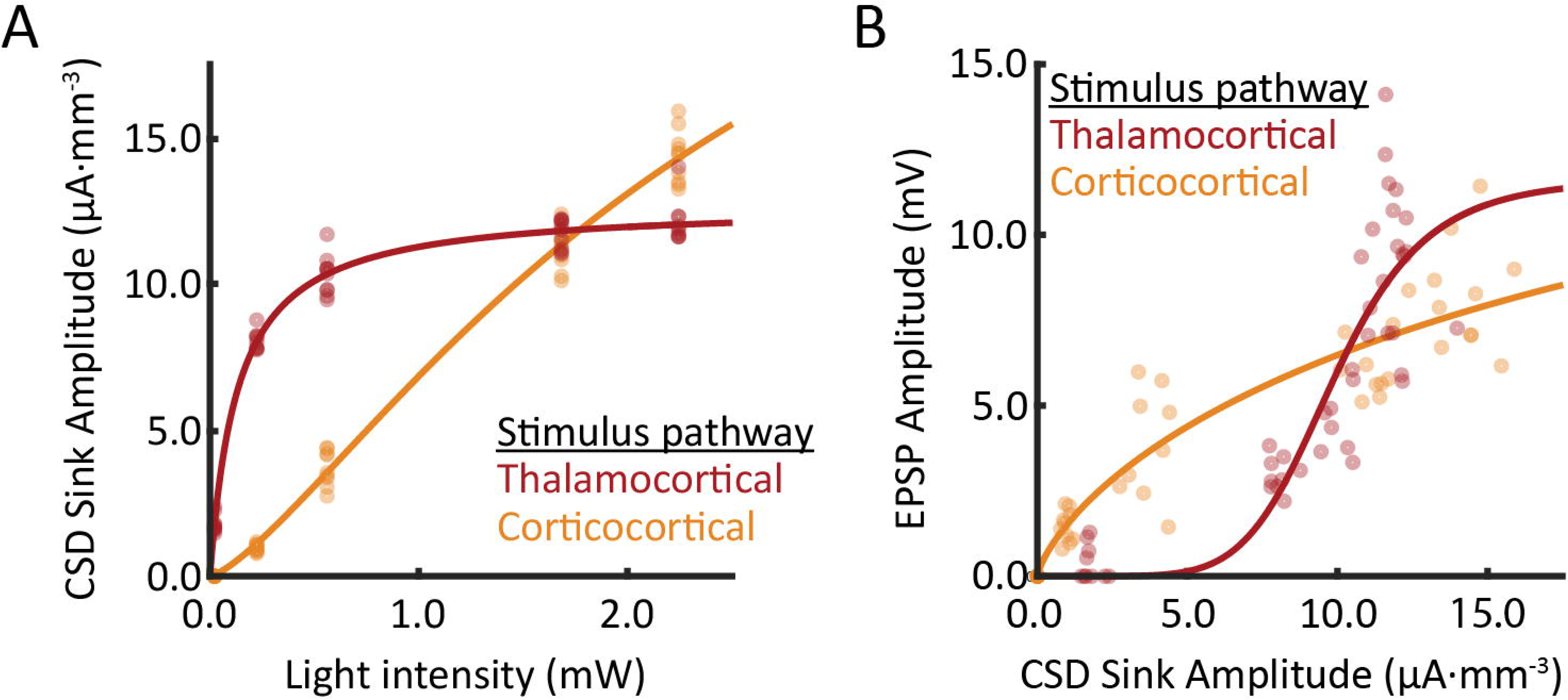
Comparison of responses from recordings in two different slices. Multiple light intensities were used to evoke synaptic responses in cortical layer 1. For each trial, the peak amplitude of the evoked response was extracted from the layer 1 extracellular current sink and EPSPs in layer 2/3 PV+ interneurons. (**A**) Extracellular response profiles of thalamocortical and corticocortical afferents are compared as a function of light intensity. (**B**) The relationship between current sink amplitude and EPSP amplitude is pathway-dependent. Within each stimulus pathway, data from (A) and (B) were collected simultaneously.

7.3. Rotate mesh grid such that the line of electrode contacts on the distal end of the multi-channel probe is approximately perpendicular to the pial surface.

7.4. Under broadfield illumination and under fine control of the micromanipulator, lower the multi-channel probe toward the surface of the slice.

7.5. Rotate the filter cube turret to engage the appropriate filter cube for visualization of the fluorescent reporter protein expressed in axon terminals of cortical afferents. If necessary, rotate the slice to more precisely align the probe with the pial surface.

7.6. Position the probe just above the plane of the slice, ∼200 µm short of the final target position along the x-axis, leaving at least one channel outside the boundary of the area of tissue being recorded

7.7. Slowly insert the probe into the slice by moving the manipulator along its longitudinal axis. To minimize damage to the tissue, only advance the probe to the extent that the sharp tips are just visible below the tissue surface. This will minimize damage to the tissue while still ensuring the electrode contacts are in contact with the tissue.

### 8. Patch clamp targeted neurons and obtain whole-cell configuration

8.1. Switch eACSF source to bagged CONTROL solution.

8.2. Identify fluorescently-labelled cell for targeted patch clamp recording.

8.2.1. Restrict the aperture iris diaphragm to the smallest diameter. Engage a low-power objective lens and bring the tissue into focus.

8.2.2. Center the light over an area of tissue adjacent to (but not overlapping) the multi-channel probe.

8.2.3. Engage the high-power (40x or 60x) water immersion objective, using caution to avoid contact between the multi-channel probe and objective lens.

8.2.4. Rotate the filter cube turret to engage the appropriate filter set to allow imaging of cells expressing Cre-dependent fluorescent marker.

8.2.5. Identify a fluorescently labelled cell as a target for patch clamp recording. Raise the objective lens to create ample space to lower a patch pipette.

8.3. Load a patch pipette (see Table 2) with internal solution (Table 1) and mount pipette into electrode holder. Using 1 mL syringe, apply positive pressure corresponding to ∼0.1mL air.

8.4. Lower patch pipette into the solution. Bring the pipette tip into focus under visual guidance.

8.5. Obtain whole-cell recording from the targeted cell using the steps previously demonstrated by Segev, et al. (2016) in their JoVE video protocol^33^.

8.6. If planning to assess changes to intrinsic properties of the cell (e.g., input resistance, action potential firing rate in response to current steps), conduct these recordings. Otherwise, move to axon stimulation protocol in below.

### 9. Layer-specific optogenetic activation of axon terminals

9.1. Manipulate field of view in the x-y plane to align light stimulation profile with desired location on slice.

9.1.1. Return the filter cube turret to the empty position. Project the image from the digital camera onto the computer screen.

9.1.2. In PolyScan2, open one of the profiles previously created in the profile sequence editor. The field of view should be just visible under the translucent Pattern Editor window from PolyScan2.

9.1.3. Under fine control, navigate to the layer of cortex that will be activated. If necessary, using the appropriate filter cube, temporarily illuminate the axon fibers by activating the viral construct fluorescent reporter. The pattern shown in the Pattern Editor should represent the exact coordinates that will be illuminated on the slice plane, visible in the digital camera’s projected image.

9.2. Engage the beam splitter in the middle position of the Mightex 3-position microscope adapter.

9.3. Load light stimulus protocol to Polygon400.

9.3.1. In PolyScan 2, close Pattern Editor and Profile Sequence Editor. Navigate to Session Control.

9.3.2. Manipulate the settings in Session Control to match those in Supplementary Figure 4. (NOTE: Make sure the value entered for PulseWidth exactly matches the Pulse width of the TTL pulse sent by the trigger within the data acquisition system.)

**Figure 4.**
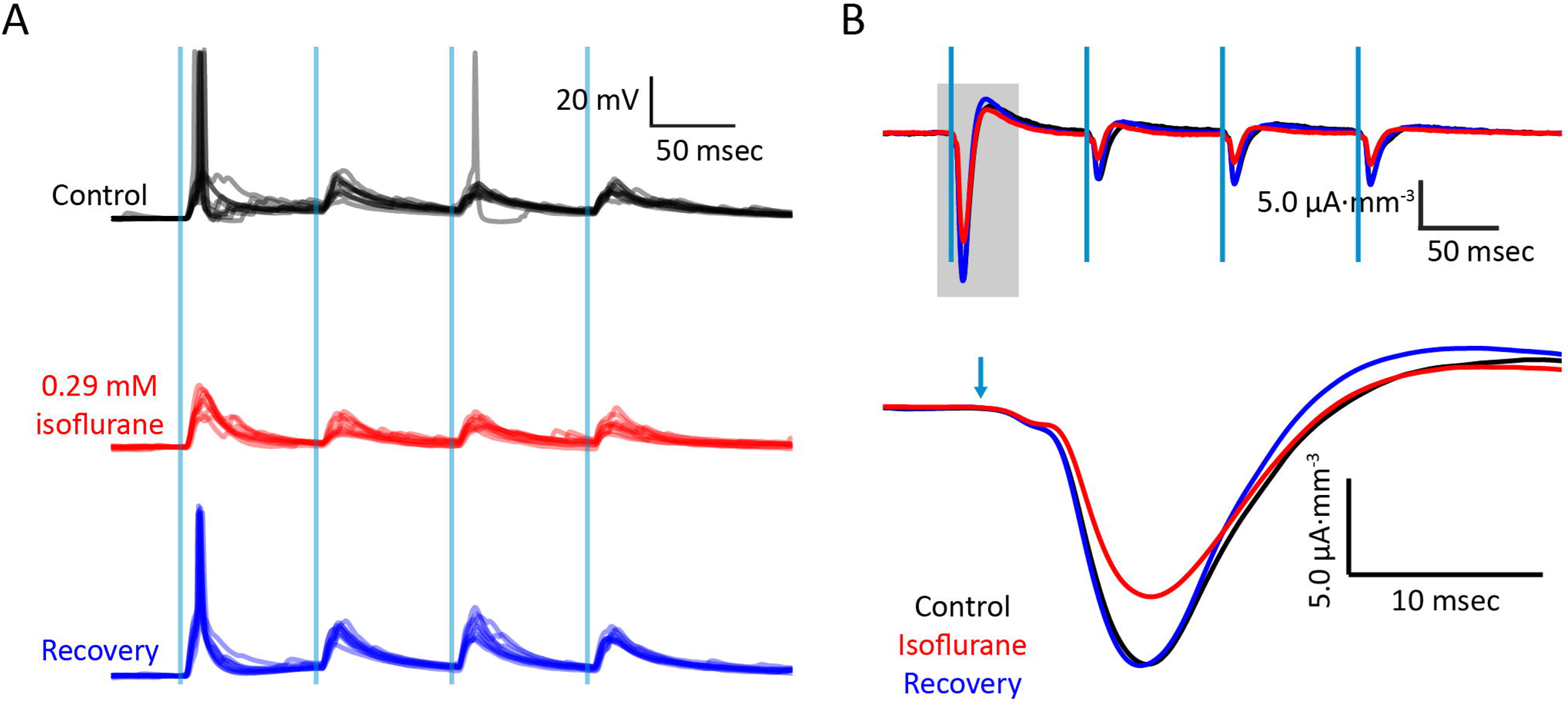
Bath application of isoflurane dissolved in eACSF during simultaneous recordings. (**A**) Intracellular whole-cell current clamp recording from layer 2/3 SOM+ interneuron upon activation of corticocortical afferents during control, isoflurane, and wash conditions. Vertical blue lines indicate light stimuli (2 msec; 1.65 mW). (**B**) Current source density trace extracted from electrode in layer 1. Data were collected simultaneously with those collected in (**A**). Recovery of responses upon wash demonstrates block of synaptic responses by isoflurane.

9.3.3. Select New. Select Add Sequence. Open the profile sequence you created in Step 6.4 above.

9.3.4. Select Start. Once PolyScan2 reads “Patterns uploaded. Polygon is ready to accept TTL triggers”, proceed to 9.4.

9.4. Optogenetically activate axon terminals while simultaneously recording extracellular field potentials and intracellular membrane fluctuations.

9.4.1. In data acquisition software, load stimulation protocol. Click record.

9.5. Following completion of all trials, click Stop in PolyScan2.

9.6. Switch eACSF source to ISOFLURANE solution and wash drug in for 15 minutes. If necessary, collect spontaneous recordings during wash-in.

9.7. Repeat steps 9.3-9.5.

9.8. Switch eACSF source to CONTROL solution and wash drug out for 20 minutes. If necessary, collect spontaneous recordings during wash-out.

9.9. Repeat steps 9.3-9.5.

## REPRESENTATIVE RESULTS

Cortical inputs arriving from higher order cortical areas or from non-primary thalamic nuclei have partially overlapping terminal fields in layer 1 of non-primary visual cortex^24^. To isolate independent thalamocortical or corticocortical afferent pathways, we injected a viral vector containing ChR2 and an eYFP fluorescent reporter (Figure 1A) into either Po or Cg. Cells within the injection radius take up the viral vector and, after 2-4 weeks, express the non-specific cation channel ChR2 and the reporter in both the soma and projecting axons. Coronal slices were collected. With the appropriate filter cube engaged, axons expressing the viral construct were imaged (Figure 1B). The use of ChR2 to activate axon terminals allows for activation of afferents without the prerequisite for an attached soma.

The animals used in the experiments described here were SOM-tdTomato or PV-tdTomato hybrid animals, which express the fluorescent reporter protein tdTomato in either somatostatin-(SOM+) or parvalbumin-positive (PV+) interneurons, respectively. SOM+ or PV+ interneurons in layer 2/3 were targeted for patch clamping under visual guidance with the appropriate filter cube engaged (Layer 1C). These interneurons have dendrites in layer 1 and are targets of corticocortical inputs (Figure 2A).

Addition of 125 mL of 3.0% isoflurane gas and 175 mL of 95% O_2_/5% CO_2_ to a sealed Teflon bag resulted in a pre-equilibrium concentration of gas of 1.3%. Gas dissolved into eACSF according to its partition coefficient; the predicted gas phase equilibrium concentration of isoflurane at room temperature was 0.6% (Figure 1D). This was confirmed via gas monitor.

The tissue slice was transferred to the recording chamber and the 16×1 multi-channel recording probe was placed orthogonally to the cortical laminae (Figure 1E). A 150 μm circle of 470 nm light centered over cortical layer 1 was delivered via the objective light path, while extracellular field potentials were collected using the 16×1 multi-channel probe and targeted whole-cell patch clamp recordings were conducted in interneurons. A schematic of the recording set-up is shown in Figure 1F.

Post-synaptic potentials (PSPs) were observed in interneurons in response to a train of four 2-msec pulses of light (10 Hz; Figure 2A). Local field potentials were also recorded (Figure 2B). Current source density (CSD; Figure 2C) and multi-unit activity (MUA; Figure 2D) were extracted from local field potentials. Ten trials at several different light intensities were used to conduct post hoc analyses. The amplitude of current sinks extracted from the CSD increased as a function of light intensity (Figure 3A). A three-parameter nonlinear logistic equation was fit to the data for comparisons across pathways. PSP amplitude also increased with current sink amplitude (Figure 3B).

Synaptic responses to thalamocortical and corticocortical inputs were measured during control, isoflurane (0.28 mM), and recovery conditions. Post-synaptic responses of somatostatin- (Figure 4A) to corticocortical stimuli were suppressed during isoflurane, as were evoked current sinks (Figure 4B).

## DISCUSSION

In this manuscript, a protocol for evaluating intra- and extracellular responses to selectively activated afferent pathways in *ex vivo* brain slices is described. The use of optogenetic tools allows investigators to probe responses of local populations to afferent inputs from distant brain regions, while recording simultaneously from targeted populations of interneurons. The use of optogenetic technology allows for axon terminals of afferent projections to be preserved and activated even though their cell bodies are no longer attached. This relieves geometric restrictions previously imposed upon *ex vivo* slices, as preservation of long-range electrical connections is no longer paramount. Still, care should be taken to prepare slices in a geometrical plane that preserves any remaining connections of interest. For example, pyramidal cells are oriented vertically along the cortical column, and evoked network activity measured by the multichannel probes in these experiments requires such local connections to be preserved as much as possible. Thus, coronal slices were prepared to keep local connectivity intact.

When choosing optogenetic constructs and relevant fluorescent reporter proteins, properties of their excitation/emission spectra and microscopic optics must be considered. Persistent light stimulation may result in partial inactivation of many channelrhodopsin variants^34^, which can be avoided by choosing reporter proteins whose excitation spectra do not overlap with that of the opsin. Alternative variants with different kinetics or light sensitivities may also be chosen depending on the experimental paradigm^35^, including manipulations using alternative excitatory or inhibitory opsins. Filter cubes must also be appropriately aligned with the chosen fluorescent reporters, such that afferent axon terminals or interneurons may be imaged independently and without activating expressed opsins.

Delivery of pre-calculated concentrations of volatile anesthetics to slice tissue is also possible using the methods outlined here. When choosing appropriate physiologically relevant gas equilibrium percentages, investigators should account for 10-15% loss of dissolved isoflurane gas between the perfusion line and tissue^36^. We have presented in detail the methods applicable to isoflurane, but other drugs such as halothane, sevoflurane, or desflurane can be handled similarly using the appropriate Ostwald and Bunsen coefficients (Supplementary Table 1). The partitioning properties of volatile anesthetics assure that they will predictably dissolve into ACSF. However, because partial pressures are more sensitive to changes in temperature than aqueous EC_50_ concentrations^31^, gas equilibrium volume percentages of volatile anesthetics must be converted to predicted room temperature millimolar concentrations to compare observed effects to physiologically relevant doses in vivo. If opting to study intravenous anesthetics such as etomidate or propofol in brain slices, investigators should consider diffusion profiles of the drugs under study, as equilibration times and physiologically relevant concentrations may vary greatly^30^.

In this manuscript, we describe a protocol for testing the effects of volatile anesthetics on distinct components of thalamocortical circuits in *ex vivo* brain slices. Many of the variables and parameters in the methods described may be manipulated for further investigations. For example, different brain areas, afferent pathways, cell targets, or volatile anesthetics may be studied by adapting the outlined methods to answer novel questions. Combined with other theoretical and experimental methods, study of unique cellular and network components using *ex vivo* brain slices will advance our understanding of the dynamic brain, and the changes it undergoes during pharmacological and pathophysiological changes in consciousness.

## FIGURE AND TABLE LEGENDS

**Supplementary Figure 1. Template for preparing block of tissue to collect brain slices.** The template is adjusted to the appropriate size, printed, and glued to a microscope slide. A cover slip should be glued over the template to prolong its use. The tissue block is placed on a piece of filter paper with the sagittal plane down, aligned to the pink background, and a vertical cut is made in the coronal plane along the black line.

**Supplementary Figure 2. Incubation chamber for harvested brain slices**. The chamber is filled with sACSF and bubbled with 95% O_2_/5% CO_2_ gas mixture via a bent needle attached to tubing. Incubation platform is made of nylon stretched over a plastic circular fitting.

**Supplementary Figure 3. Platinum structures for slice in recording chamber**. Brain slice is transferred to recording chamber via pipette and placed on top of nylon mesh, which is stretched over a horseshoe-shaped piece of flattened platinum wire and super glued in place. Platinum harp is placed over brain slice to anchor it in place during recording.

**Supplementary Figure 4. Screenshot of PolyScan2 software prior to recording**. Settings should be toggled as shown here to prepare Polygon400 to receive TTL input from digital recording software. From this window, a new stimulation protocol is chosen and uploaded to Polygon400.

**Supplementary Table 1. Ostwald (λ) and Bunsen (α) coefficients for other volatile anesthetics**. This protocol may be adapted for study of other volatile gas anesthetics, such as halothane, sevoflurane, or desflurane. The equations described in the protocol should be substituted with the appropriate coefficients as listed in this table.

## ACKNOWLEDGMENTS

This work was supported by the International Anesthesia Research Society (IMRA to AR), National Institutes of Health (R01 GM109086 to MIB), and the Department of Anesthesiology, School of Medicine and Public Health, University of Wisconsin, Madison, WI, USA.

---

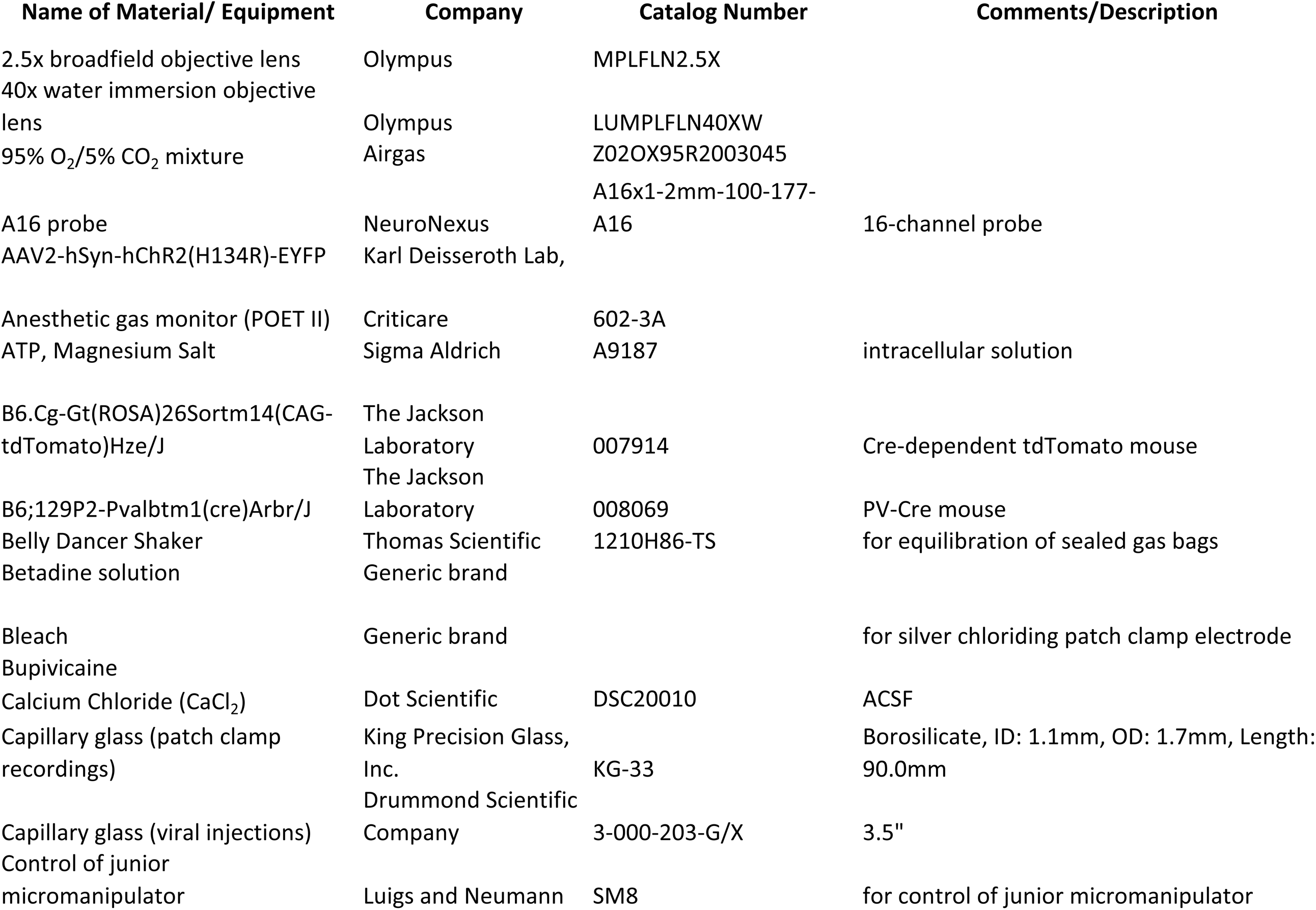
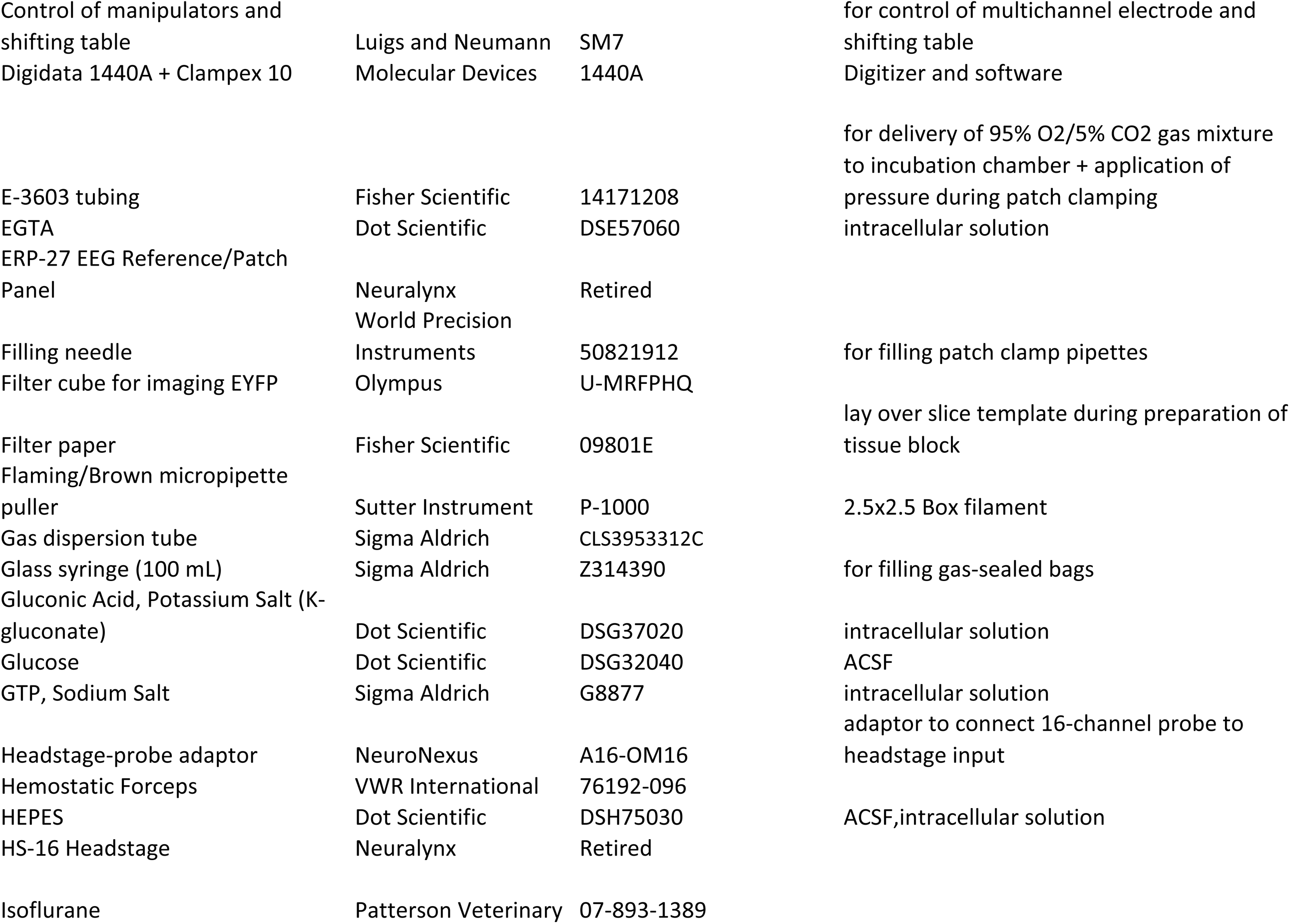
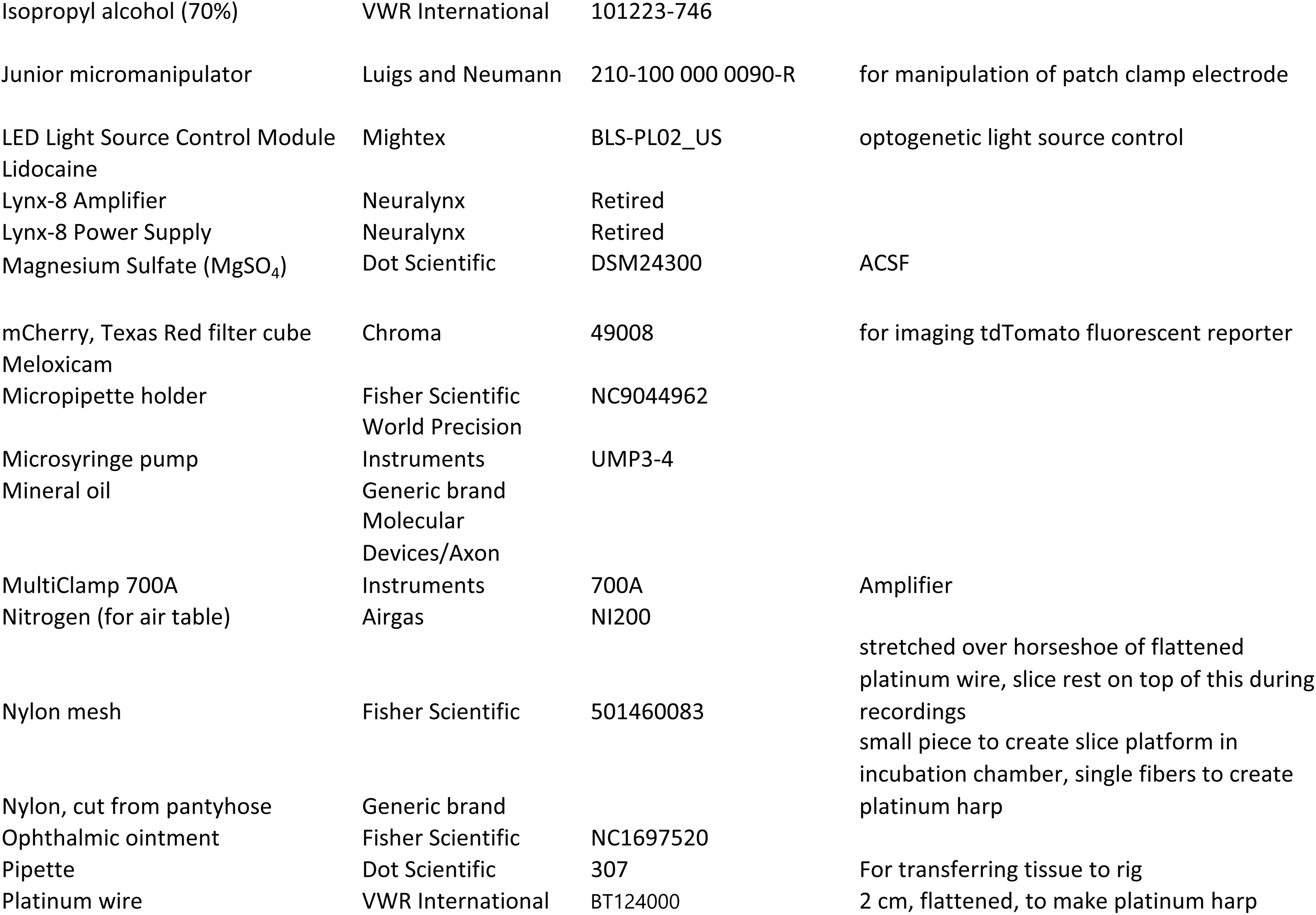
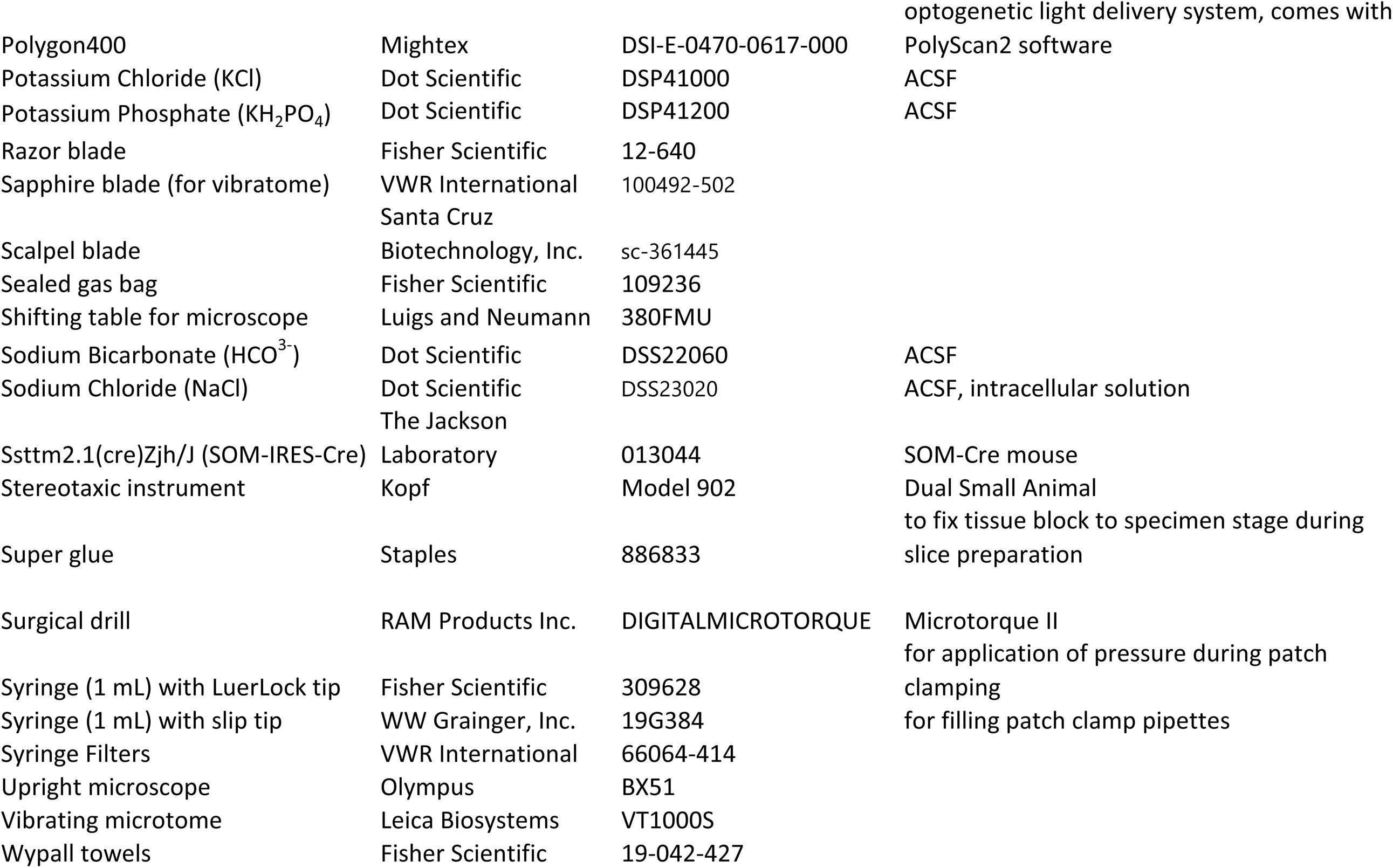

