## Supplementary figures and images for "Optogenetic activation of afferent pathways in brain slices and modulation of responses by volatile anesthetics"

### Supplementary Figure 1

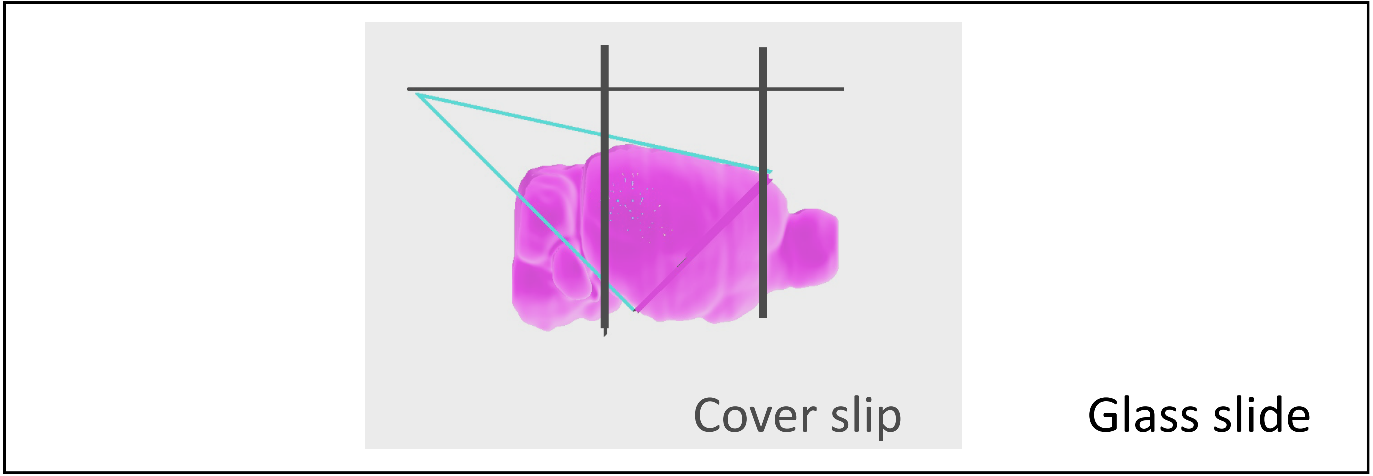

### Supplementary Figure 2

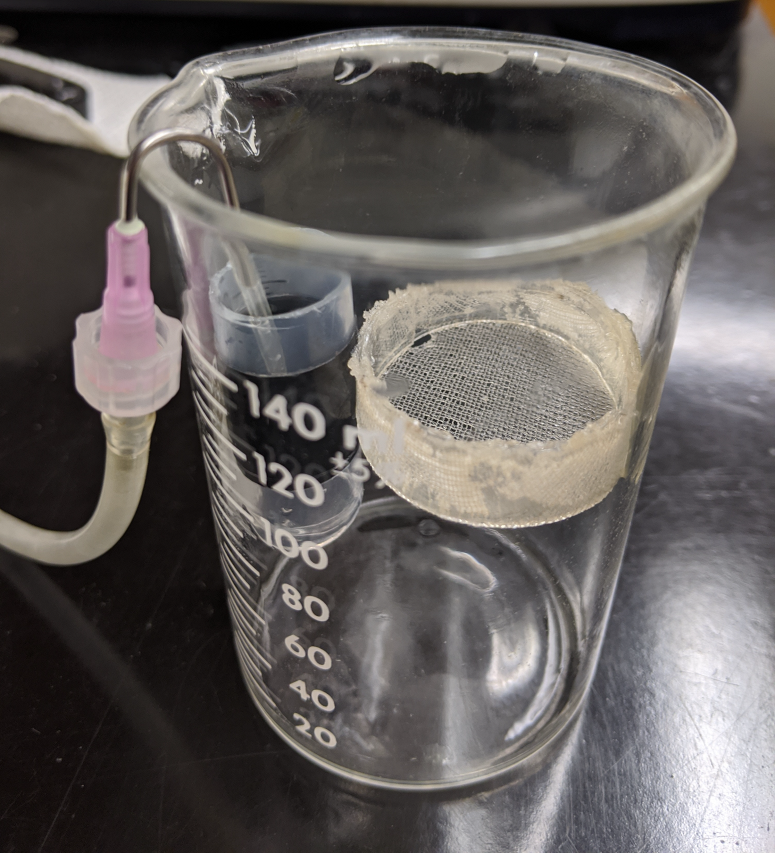

### Supplementary Figure 3

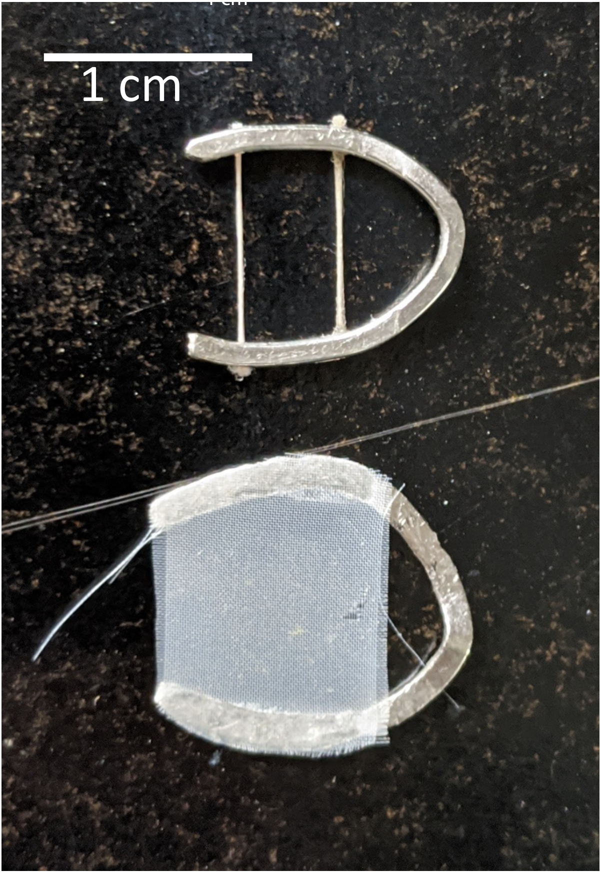

### Supplementary Figure 4

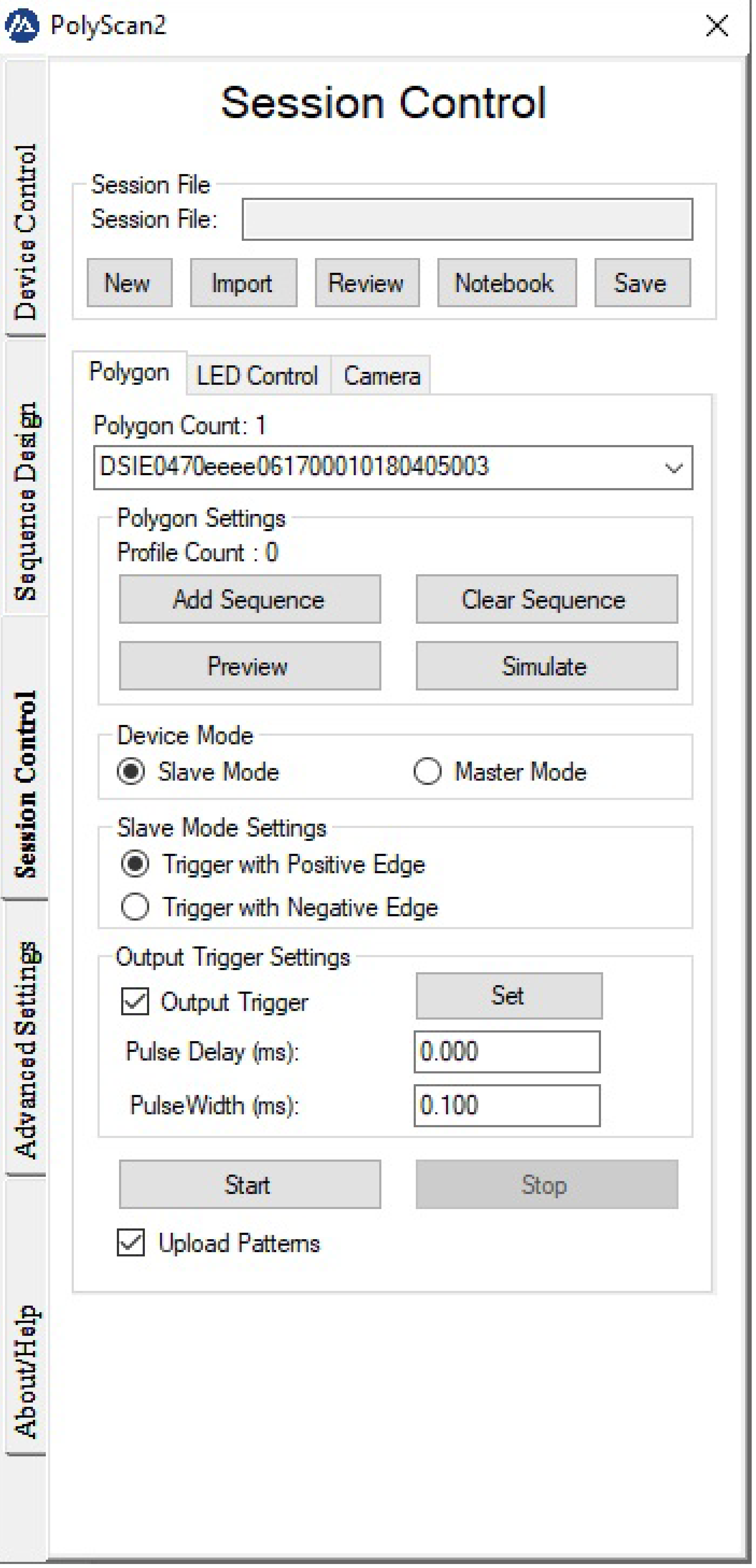
